# Heuristic hyperparameter optimization of deep learning models for genomic prediction

**DOI:** 10.1101/2020.11.25.398800

**Authors:** Junjie Han, Cedric Gondro, Kenneth Reid, Juan P. Steibel

**Affiliations:** Department of Animal Science, Michigan State University, East Lansing, MI 48824; Department of Computational Mathematics, Science and Engineering, Michigan State University, East Lansing, MI 48824; Department of Fisheries and Wildlife, Michigan State University, East Lansing, MI 48824

**Keywords:** Genomic Prediction, Deep Learning, Hyperparameter optimization, Evolutionary Algorithm

## Abstract

There is a growing interest among quantitative geneticists and animal breeders in the use of deep learning (DL) for genomic prediction. However, the performance of DL is affected by hyperparameters that are typically manually set by users. These hyperparameters do not simply specify the architecture of the model, they are also critical for the efficacy of the optimization and model fitting process. To date, most DL approaches used for genomic prediction have concentrated on identifying suitable hyperparameters by exploring discrete options from a subset of the hyperparameter space. Enlarging the hyperparameter optimization search space with continuous hyperparameters is a daunting combinatorial problem. To deal with this problem, we propose using differential evolution (DE) to perform an efficient search of arbitrarily complex hyperparameter spaces in DL models and we apply this to the specific case of genomic prediction of livestock phenotypes. This approach was evaluated on two pig and cattle datasets with real genotypes and simulated phenotypes (N=7,539 animals and M=48,541 markers) and one real dataset (N=910 individuals and M=28,916 markers). Hyperparameters were evaluated using cross validation. We compared the predictive performance of DL models using hyperparameters optimized by DE against DL models with “best practice” hyperparameters selected from published studies and baseline DL models with randomly specified hyperparameters. Optimized models using DE showed clear improvement in predictive performance across all three datasets.DE optimized hyperparameters also resulted in DL models with less overfitting and less variation in predictive performance over repeated retraining compared to non-optimized DL models.

## Introduction

Over the past decades, there have been enormous gains in the productivity of livestock, much of which was due to the rapid genetic improvement of quantitative traits e.g. growth rates, reproductive traits, and feed conversion rates (Hill 2016). In recent years, with the rise of DNA sequencing and high throughput genotyping technology as well as with the inception of genomic prediction models (Meuwissen *et al.* 2001), single nucleotide polymorphisms (SNP) became widely used for genomic prediction and genomic selection.

Genomic prediction refers to the use of statistical models to estimate the genetic component of a phenotype by using data from thousands of SNP markers (Meuwissen *et al.* 2001; Pérez-Enciso and Zingaretti 2019). The same models can also be used for phenotypic (genomic) prediction by associating an individual’s genotype to its phenotypes which is commonly used to predict complex traits in humans (Yang *et al.* 2010). For animal production, both genomic prediction and phenotypic prediction have resulted in faster and more accurate selection while genomic prediction has been useful for management decisions (e.g. market allocation). The technology has also provided a platform for the adoption of novel breeding approaches and has led to new biological insights into the underpinnings of complex quantitative traits (Hickey *et al.* 2017). For simplicity we will use only the term genomic prediction throughout the text.

Several models have been proposed for genomic prediction (VanRaden 2008; Corvin *et al.* 2010; Habier *et al.* 2011; Gianola 2013), and GBLUP is one of the most commonly used models (Fragomeni *et al.* 2017). A common assumption across these models is that genomic effects are strictly additive, i.e. most models do not explicitly consider interactions between alleles within markers (dominance), nor between markers (epistasis) (Crossa *et al.* 2019). More recently, deep learning (Lecun *et al.* 2015) has been proposed as an alternative to genomic prediction models that does not depend on the typical assumptions of traditional genomic prediction methods.

Deep learning (DL) has dramatically improved state-of-the-art applications in computer vision, speech recognition and genomics (Lecun *et al.* 2015). DL methods are flexible and can potentially learn very cryptic data structures - even interactions between predictors (Crossa *et al.* 2019). DL has already been applied to genomic prediction in plants (Montesinos-López *et al.* 2018; Crossa *et al.* 2019), human traits (Bellot *et al.* 2018), and estimation of breeding values in cattle (Abdollahi-Arpanahi *et al.* 2020).

DL models in genomic prediction are promising tools (Bellot *et al.* 2018). However, one of the critical challenges of implementing DL is selection of appropriate hyperparameters since they significantly affect the performance of the prediction algorithm. Hyperparameter features are values or options typically set by users before the model is fitted that impact the algorithm’s predictive performance by avoiding overfitting and underfitting (Luo 2016). Each feature that is part of the hyperparameter set can take a range of values or options and they can interact with each other to determine the properties of the final fitted model; a properly specified hyperparameter set is fundamental for a DL model to achieve a high prediction accuracy. but, unfortunately, there is no formalized or standard way to optimize these hyperparameters.

Several procedures have been used to select DL hyperparameters for genomic prediction applications; e.g. grid search (Crossa *et al.* 2019; Pérez-Enciso and Zingaretti 2019) and genetic algorithms (Bellot *et al.* 2018). Grid search is only feasible for a limited number of parameters and levels, which is not the case for most DL applications. On the other side, genetic algorithms are better suited for optimizing large and complex parametric spaces, but currently available implementations of genetic algorithms to tune DL hyperparameters for genomic prediction require that the options of each hyperparameter are either already discrete or discretized before the optimization process (Bellot *et al.* 2018).

An alternative to genetic algorithms is differential evolution (DE) which is a population based evolutionary heuristic well suited for optimization of discrete and continuous search spaces (Storn and Price 1997; Das *et al.* 2016). Differential Evolution lies on the intersection between real-valued genetic algorithms and evolution strategies. DE uses the conventional population structure of genetic algorithms and the self-adapting mutation of evolution strategies; in a sense DE can be loosely viewed as a population based simulated annealing algorithm in which the mutation rate decreases as the population converges on a solution.

In this study, we propose how to adapt DE to optimize the DL hyperparameter set for genomic prediction and evaluate its effectiveness to improve prediction accuracies in simulated and real datasets for two classes of DL models: multilayer perceptron (MLP) and convolutional neural network (CNN). We emphasize that the focus of this paper is on optimization of DL hyperparameters to identify a set suitable for a given specific genomic prediction problem, rather than a comparison of DL with GBLUP as a genomic prediction method. As the predictive performance depends on the architecture of the trait and the population structure, we demonstrate the importance and the impact of proper hyperparameter specification on genomic prediction with DL.

## Material and Methods

### Simulated datasets

Real genotypes from two livestock populations - pigs and cattle - were used to create simulated datasets for testing purposes. Genotypes from both species were edited to be of the same dimensions, comprising a total of 48,541 SNP genotypes for 7,359 individuals, from which 6,031 (80%) and 1,508 (20%) were randomly assigned to the discovery and validation populations, respectively. Phenotypes were simulated for both species by randomly assigning 1,000 SNP as quantitative trait loci (QTL) with additive effects for a heritability of 0.4 using the R simulation package GenEval (Cuyabano 2020; R Core Team 2020).

### Real dataset

The real data came from an experimental F2 cross of Duroc and Pietrain pigs already previously described (Edwards *et al.* 2008). Briefly, four Duroc sires were mated to 15 Pietrain dams to produce 56 F1 individuals (50 females and 6 males). F1 animals were mated to produce a total of 954 F2 pigs that were phenotyped for 38 meat quality and carcass quality traits. For this study pH meat records measured 24 hours post-mortem from 910 F2 pigs were used. We purposely selected this trait as it is moderately heritable (h^2^=0.19±0.05) and for which we have mapped putative QTL (Casiró *et al.* 2017). Two different SNP chips were used to genotype the F2 pigs, but all SNP were imputed to a common set of approximately 62,000 SNP (Gualdrón Duarte *et al.* 2013) with high accuracy (R2>0.97). SNP were pruned by filtering out SNP with: 1) low genotyping rates, 2) lack of segregation, 3) inconsistent Mendelian inheritance with the pedigree information, 4) low imputation accuracy (Casiró *et al.* 2017), 5) and high correlation between markers (larger than 0.99). A final set of 28,916 SNP was used for this study.

Phenotypic records were pre-adjusted for fixed effects:

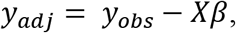

where *y_adj_* is adjusted response, *y_obs_* is pH measured 24 hours post-mortem, *X* is the incidence matrix with the fixed effects of sex, slaughter group and carcass weight, and *β* represents the coefficients of fixed effects.

### Deep learning and genomic prediction

Deep learning (DL) methods are a set of representation learning methods, where a machine can be fed with raw data and automatically discover the representations needed for prediction or classification, with multiple levels of simple but non-linear modules that transform the representation at one level into a representation at a higher, slightly more abstract level (Lecun *et al.* 2015). In the context of genomic prediction, we used DL to build a system that predicts an animal’s phenotypic value given its genotype. DL computes and minimizes a loss function that measures the error of prediction. In this study, we used mean squared error (mse) as the loss function:

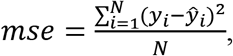

where *N* represents the number of individuals in the training dataset, *y_i_* represents the observed response of individual *i* and *ŷ_i_* is the predicted response of individual *i*. Two types of DL models were used in this study: multilayer perceptron and convolutional neural network.

### Multilayer perceptron

This model is also known as feed-forward artificial neural network. In this paper, MLP (Figure 1) has: an input layer with as many nodes as SNP markers, a variable number of hidden layer(s) with a certain number of nodes, and an output layer representing the response. Since nodes between layers are fully connected, MLP can potentially model complex and higher order interactions between predictor variables (Abdollahi-Arpanahi *et al.* 2020). A detailed explanation of how MLP models work is presented in the File S1 and also available at GitHub alongside source code (https://github.com/jun-jieh/DE_DL).

**Figure 1.**
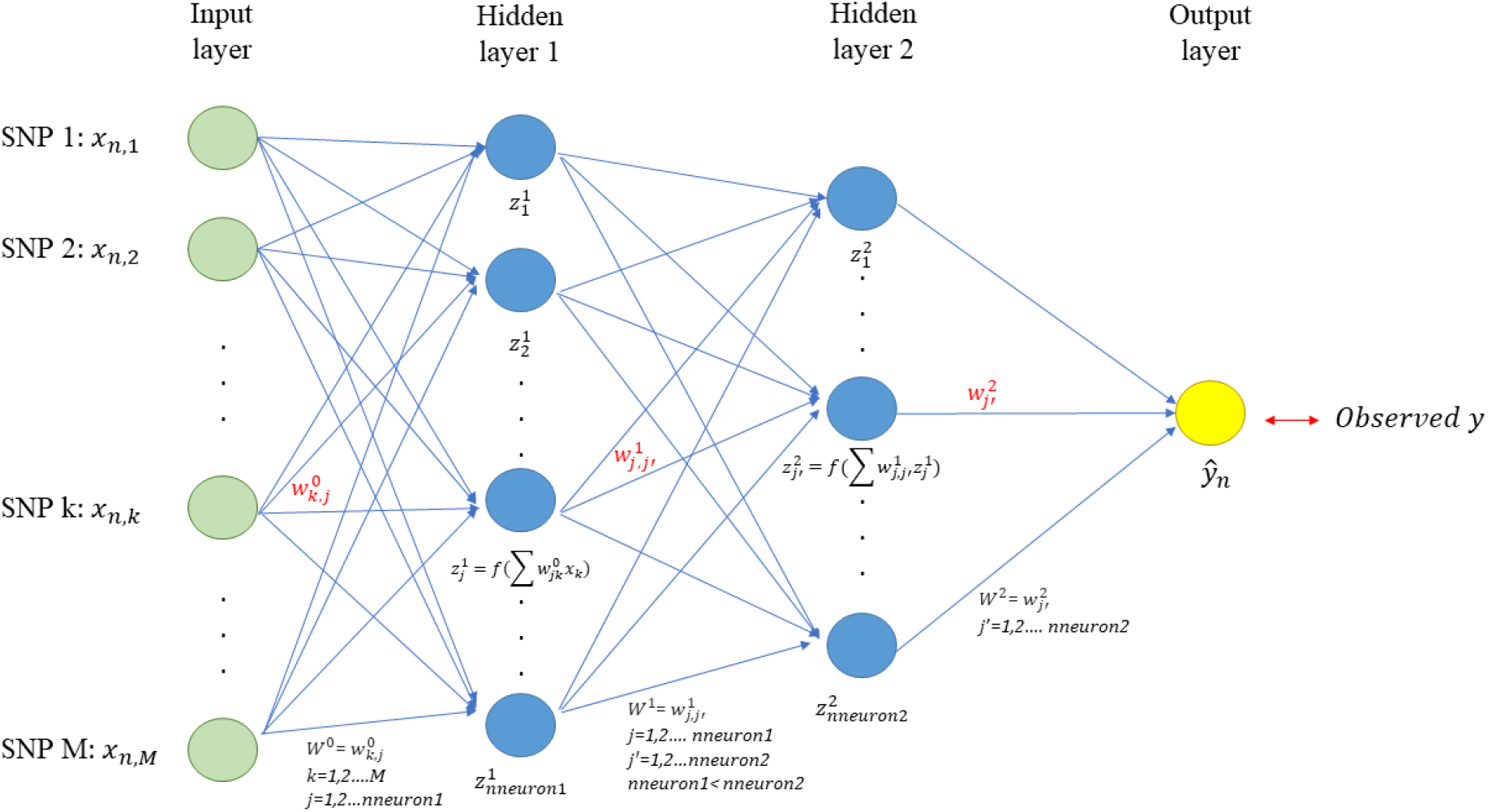
Multilayer Perceptron (MLP) for genomic prediction of a single trait with M SNP markers. The network has an input layer, two fully connected hidden layers and an output layer. Each node’s input in the hidden layers is a transformation of the weighted sum of the output from the previous layer. The number of nodes in hidden layers decrease as the depth of the MLP increases, to facilitate representation learning.

As deep learning consists of transforming representations at a previous layer into its next (more abstract) layer (Lecun *et al.* 2015), we opted to adaptively set the number of nodes for each hidden layer based on the depth of the network so that the next hidden layer always has fewer nodes than the previous one. For instance, in an MLP with two hidden layers (Figure 1), the first layer can only have neurons ranging from 259 to 512 while the second layer can have any number between 4 and 258. Table S1 summarizes the number of nodes search space for MLPs with one, two, and up to five layers. Other researchers may choose different adaptive rules to impose restrictions on the possible number of neurons per layer, or may even simply choose to use the same number of nodes for all layers.

### Convolutional neural network

CNN is designed to process data that comes as multiple-array format (Lecun *et al.* 2015) e.g. 1d for an animal’s genotype, 2d for images and 3d for videos. Typical CNN models consist of an input layer, convolutional layer(s), pooling layer(s), a flattened layer, and an output layer (Figure 2). In the context of genomic prediction (Figure 2), the input layer for a single observation in a CNN is a one-dimension array that contains an animal’s genotype and the number of units in the layer will be equal to the number of markers. The output layer *ŷ_n_* represents the predicted response value for the phenotype or breeding value of the *n^th^* individual.

**Figure 2.**
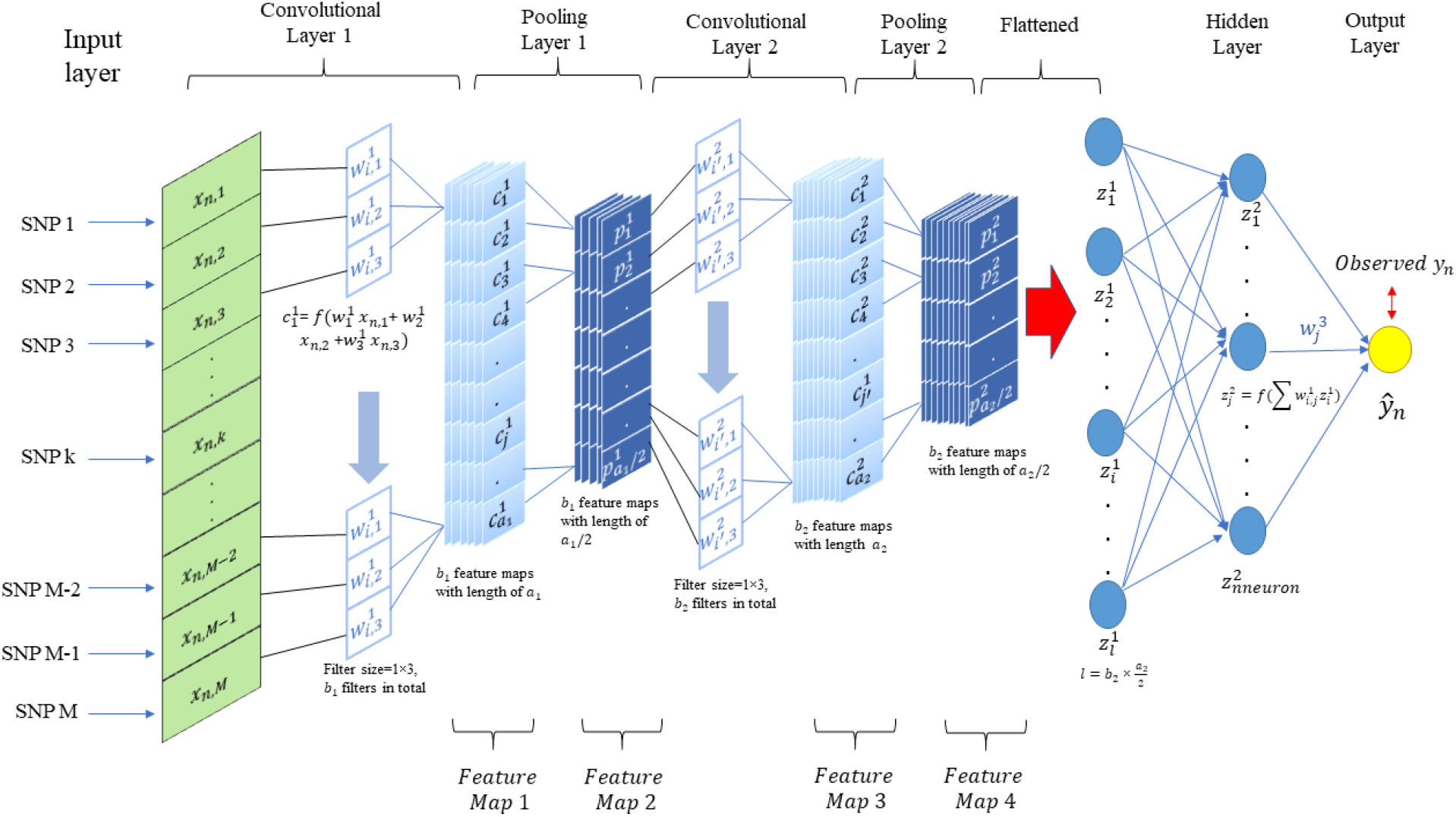
1-d Convolutional neural network (CNN) for genomic prediction of a single trait with M SNP markers. The network has an input layer, two convolutional layers with their corresponding pooling layers, a fully connected hidden layer and an output layer. Each convolutional layer applies a number of filters to the output of the previous layer and its output is subsequently summarized by a pooling layer, where filters are arrays for convolving input. The number of filters generally increase as the network becomes deeper and each filter learns a different abstract representation of the input data from a previous layer.

Between the input and output layers, a CNN contains a variable number of convolutional layer(s) followed by pooling layer(s). Full details on CNN architecture are given in the File S1 and at GitHub along with source code (https://github.com/jun-jieh/DE_DL). In this study each convolutional layer applied filters of size *F* (a hyperparameter to be optimized) with the stride equal to the filter size (non-overlapping convolutions of the input). In CNN, several restrictions are typically assumed regarding the model architecture. When learning from a global level to a local level, more details are required to obtain the pattern at the local level (Lecun et al., 2015). Therefore, the number of filters increases as the depth of the CNN increases, to detect local motifs. To reflect this expectation we adaptatively set the number of filters applied in each convolutional layer as a function on the depth of the network. Specifically, we limited the number of filters in any convolutional layer to be between 4 and 128, but this range is partitioned for each convolutional layer to make sure that the next convolutional layer will always have a number of filters larger than the previous layer. For example, in a CNN with two convolutional layers (Figure 2), the first convolutional layer can only have between 4 and 65 filters while the second convolutional layer can have between 66 to 128 filters. Examples of the adaptive number of filters as a function of the depth of the CNN is presented in Table S2. The hyperparameter space for filter size was set as an integer between 2 and 20. Although the filter size is specified by the user, the output feature (feature map in Figure 2) has to conform to the minimum length (the length of feature map needs to be equal to or larger than the filter size) of the feature maps in each convolutional layer and pooling layer, which is illustrated in Table S3. If the condition is not satisfied, instead of fixing the kernel size through all convolutional layers, we set adaptive kernel size in order to successfully execute the model fitting. The adaptive kernel size is to ensure that CNN generates a valid output.

### DL model training

TensorFlow (Abadi *et al.* 2015) was used to train DL models. Noteworthy, at each iteration (epoch) of the training process TensorFlow randomly partitioned the training data into an actual training set, that was used for updating the model weights and an internal validation set, that was used to compute the correlation between predicted and observed response (a detailed description of epoch is presented in File S1). A DL training procedure typically requires multiple epochs, which was one of the hyperparameters optimized in our DE procedure (see below).We also introduced an early stopping when the correlation did not change over 0.1 for ten consecutive epochs, as it was assumed that fitness could not be improved and further exploration was unnecessary.

### Hyperparameter optimization

Table 1 presents the hyperparameters optimized in this study. A plausible range of values for each hyperparameter was defined based on ranges suggested by the literature for DL applied to genomic prediction (additional details of each hyperparameter can be found in the File S1). These ranges were then used as constraints for the differential evolution algorithm.

**Table 1.** Parameter space for optimized hyperparameters

| Hyperparameters | Parameter space (MLP) | Parameter space (CNN) | Value Type |
| --- | --- | --- | --- |
| Number of layers | [1,2,3,4,5] | [1,2,3,4,5] | Integer |
| Number of neurons | [8-512] | [8-512] | Integer |
| Activation | ['relu', 'elu', 'sigmoid', 'selu', 'softplus', 'linear', 'tanh'] | ['relu', 'elu', 'sigmoid', 'selu', 'softplus', 'linear', 'tanh'] | Categorical |
| Optimizer | ['sgd', 'adam', 'adagrad', 'rmsprop', 'adadelata', 'adamax', 'nadam'] | ['sgd', 'adam', 'adagrad', 'rmsprop', 'adadelata', 'adamax', 'nadam'] | Categorical |
| Dropout rate | [0-1] | [0-1] | Continuous |
| L2 penalty | [0-1] | [0-1] | Continuous |
| Batch size | $[N \times \alpha_1 - N \times \alpha_2]$ | 32 | Integer |
| Epoch | [21-50] | [21-50] | Integer |
| Number of filters | NA | [2-128] | Integer |
| Filter size | NA | [2-20] | Integer |
| Pooling | NA | ['max', 'average'] | Categorical |

Hyperparameter space and range (see details in File S1). N represents sample size. $\alpha_1=0.001$ for the simulated datasets and $\alpha_1=0.01$ for the real pig dataset. $\alpha_2=0.01$ for the simulated datasets and $\alpha_2=0.1$ for the real pig dataset.

### Differential evolution algorithm for deep learning

Differential evolution (DE) is an evolutionary algorithm that includes four steps: 1) initialization, 2) mutation, 3) crossover and 4) selection (Storn and Price 1997). A generic version of this algorithm is described in pseudocode format (Figure 3). DE was used to evolve a population of numeric vectors that can be recoded to represent hyperparameter combinations through random keys.

**Figure 3.**
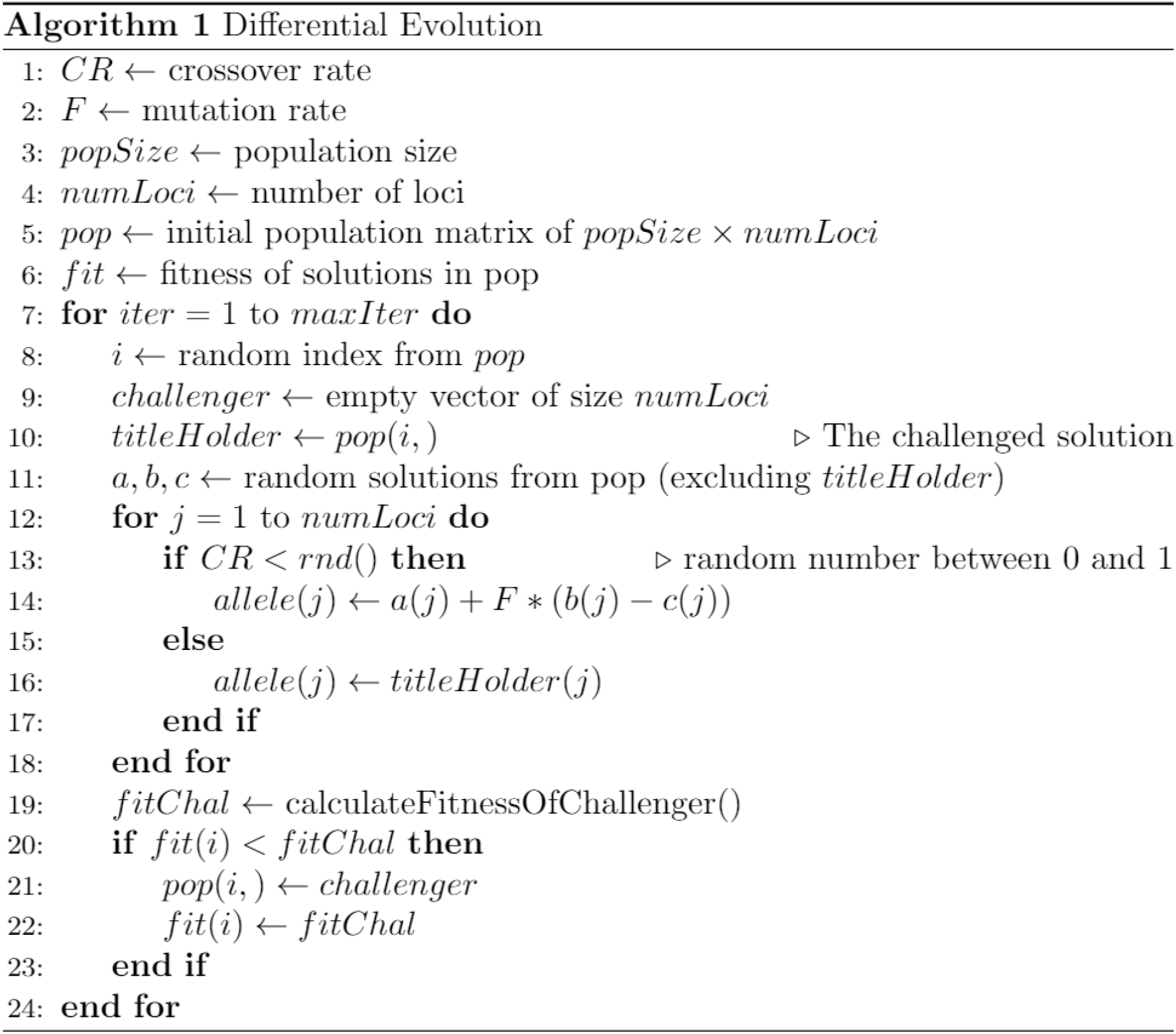
Pseudocode for differential evolution algorithm

### Random key

Random key is an encoding mechanism originally used in genetic algorithms by Bean (1994). The core of this algorithm is a set of *d H*-dimensional numeric vectors (individuals) ***pop_1_*, ..., *pop_d_*** as a population. Each numeric vector represents an individual that is linked or mapped to a set of model hyperparameters through a mapping function (random key). Suppose that there are *K* hyperparameters to optimize, where *K=8* for the MLP and *K*=10 for the CNN (Table 1). Within each hyperparameter *k*=1...*K* there are *H_k_* loci, and if the parameter takes continuous values, then *H_k_* =1. So, the size 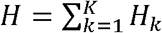. Each vector *pop_i_* is partitioned into *K* sub-blocks that contain *H_k_* loci, where each single locus in *H* represents a hyperparameter option or value. For categorical hyperparameters, there is a mapping performed from the *H_k_* dimensional block of the numeric vector to an *H_k_* dimensional vector MAP_k_ containing the names of the categories for the *k^th^* hyperparameter as follows: the *H_k_* elements are ranked according to their values and the rank of the first element is used as an index for the MAP_k_ vector to select the corresponding categorical value. In this way, the evolutionary operators (mutation, crossover, and selection) can be applied directly on the numerical vector *pop_i_* but the results can always still be translated into a set of categorical (and continuous) hyperparameter values. An example of this with the hyperparameter number of layers (*H_k_* =5) is presented in Figure 4. So, in a nutshell, a random key is a vector of real numbers that, once sorted, its ranking can be used to map against a set of statically ordered features. The idea is that better features will evolve to higher values in the key while worse features will evolve to lower values; the ranking of the sorted key allows sorting the features from best to worst and provides a smoother fitness surface for the DE to explore.

**Figure 4.**
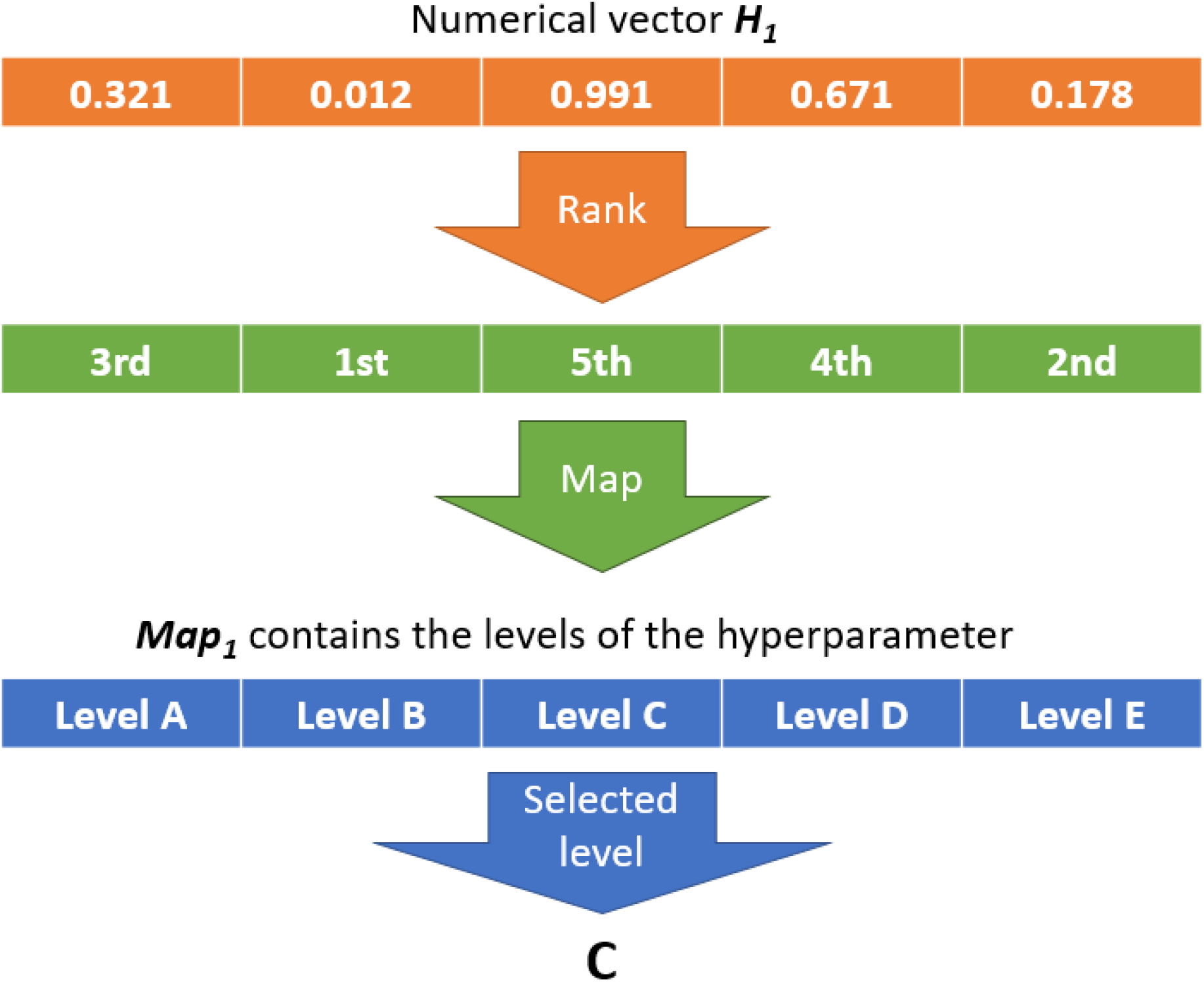
Summary of the random key (mapping function) used to transform numeric vectors into discrete levels of hyperparameters. The numeric vector can be subject to mutation and recombination. The mapping is used to transform the result into a meaningful set of hyperparameters that can be used to fit a model and obtain a fitness to select numeric vectors.

The main steps for the differential evolution algorithm are:

1. **Initialization** We initialized *d*=50 *H*-dimensional parameter vectors *pop_1_.,.pop_50_* as a population *pop* (line 5 Figure 3) from a uniform [0,1] distribution and we mapped the numeric vector to a set of hyperparameter values as described before (Table 1) to obtain 50 hyperparameter sets. Then we fitted 50 models using each set of hyperparameters and recorded their correlations between predicted and observed response values.
2. **Mutation** To generate a mutation, indexes of two random individuals are selected from the population: r_1_, r_2_, ∈ {1,2, ...,*d*} and the target *H*-dimensional vectors *pop_r_1__* and *pop_r_2__* are extracted. Then vectors *p* and *pop_r_2__* are mutated using

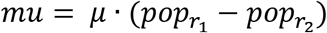

where *mu* is the mutant vector and *μ* is the mutation parameter (*μ* ∈ [0,1]). Storn and Price (1997) recommended that 0.5 is usually a good initial choice for the mutation. In this study, we set *μ* = 0.5.
3. **Crossover** To increase the diversity in hyperparameter combinations represented in the population parameters, crossover function is used to combine the mutant vector *mu* with other individual vectors. First, an *H*-dimensional vector *RN* with uniformly random numbers ∈ [0,1] is generated. The crossover rate is defined by parameter *α (α ∈* [0,1]). Gämperle et al. (2002) suggested that a good choice for the crossover constant is a value between 0.3 and 0.9. In this study, we set *α =* 0.5. Another *H*-dimensional vector (*CR*) with logical variables (True/False or 1/0) is then generated according to

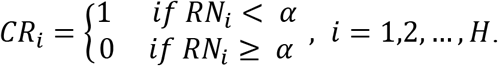 Then, two more individual vectors *pop_r_3__* and *pop_r_4__* are selected and crossover generates a new individual according to

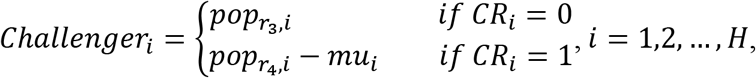

where *Challenger* is the newly generated individual and *i* is the *i^th^* element of *pop_r_3__*, *pop_r_4__*, *mu*, and *Challenger*. For convenience, we name *pop_r_3__* Titleholder.
4. **Selection** To decide whether or not the *Challenger* should replace the *Titleholder* in population *pop,* both vectors of numeric values *Challenger* and *Titleholder* were mapped into hyperparameter sets using the random key. The models were fitted based on mapped hyperparameters and Pearson correlation coefficient γ between the predicted and the observed values was computed. We averaged the correlations over epochs, as the internal validation sets varied in epochs. The averaged correlation was defined as the fitness of the DL model (given a hyperparameter set). Additionally, we applied a penalized fitness if any of the following three scenarios happened during model fitting: exhausted memory when fitting a specified model, a constant generated for all predicted responses, and exploding/vanishing gradient which led to an unstable model-fitting procedure (convergence issue). A penalized individual had its fitness set to −1. If *γ_Titleholder_ < γ_challenger_*, we replaced *Titleholder* by *Challenger* in *pop*; otherwise, we retained *Titleholder* in *pop.* Finally, steps 2)- 4) are repeated for *δ* iterations. For the simulated pig dataset and the simulated cattle dataset, *δ* of both datasets was 2,000, while *δ* = 10,000 was used for the real pig dataset (after *δ* iterations there was no further significant improvement). It is worth noting that the initial population does not need to be random, it can be based on prior information or can even be the results from a previous run -in effect, the DE can continue evolving a population that has already optimized for some iterations if e.g. the run did not converge.

As shown in the results section, if a DL model is run multiple times with the same dataset and hyperparameters, the predictive performance differs slightly from run to run. This means that a model trained once can get a slightly higher/lower prediction accuracy compared to the average prediction accuracy that would be obtained over multiple re-trainings. This effect is more pronounced in more complex models, which are more prone overfitting. To mitigate this problem, we introduced a variation to the traditional DE algorithm by refitting the *Titleholder* each time and updating its fitness value. Specifically, in each iteration, the *Titleholder* was refitted, and if the *Titleholder* won the contest, the updated fitness was retained.

### Top model selection

At the end of the DE run, each individual solution in the population was refitted 30 times to select the best model based on two criteria to evaluate model stability through repeated training of each DL model. The best model was selected based on two measures obtained from this repeated training: mean fitness and standard deviation (SD) of the fitness obtained by refitting each model 30 times. This is necessary because as explained above, the refitting of the selected models resulted in slightly different predictive performance. The details on how this bi-variate criteria selection was performed can be found in GitHub (https://github.com/jun-jieh/DE_DL).

### Optimized model assessment through external validation

Each dataset was partitioned into five training sets and five validation sets (80% and 20% for training and validation, respectively). The DE was run on each of these training sets to optimize hyperparameter sets for both, MLPs and CNNs. The final MLP and CNN models (2×5) from the DE runs were then refitted 30 times (with the training dataset only) and each refitted model was evaluated by predicting the corresponding validation set and computing the correlation between the predicted and the observed response. The average correlation for each of the 30 refits as well as the SD of correlations was then calculated. This external cross validation is distinct from the internal cross validation utilized by the DE to optimize the hyperparameters (described in the DL model training section) and should be differentiated. GBLUP was used to estimate the response variable and its prediction accuracy as a comparison reference to the optimized MLPs and CNNs (GBLUP details can be found in File S1).

### Hardware and software

The computer processor used in this study was Intel(R) Core i7-8750H CPU @ 2.20 GHz with 16GB of RAM memory and Microsoft(R) Windows 10 operating system. The graphic card was NVIDIA(R) GeForce GTX 1070 with 8 GB GDDR5 memory. All the analyses were implemented in R (R Core Team, 2020). For GBLUP we used the *gwaR* R package (Steibel 2015) and for DL the R *Keras* package (Chollet 2017), which is a high-level neural networks API on top of *TensorFlow* (Abadi *et al.* 2015).

### Data availability

The authors state that all data necessary for confirming the conclusions presented in the article are represented fully within the article. Animal protocols were approved by the Michigan State University All University Committee on Animal Use and Care (AUF# 09/03-114-00). Custom R code used to fit MLP/CNN, implement DE, and evaluate models are available at GitHub (https://github.com/jun-jieh/DE_DL). Genotypes and phenotypes for the animals in the real pig dataset are available at GitHub (https://github.com/jun-jieh/RealPigData).

## Results and discussion

As reported in many published applications of deep learning in genomic predictions (Bellot *et al.* 2018; Abdollahi-Arpanahi *et al.* 2020; Zingaretti *et al.* 2020), we observed that retraining of a certain DL model with the same hyperparameter configuration and the same dataset produced slightly different predictions. This forced us to consider the variation in the predictive performance under the retraining in DE and post-DE model selection (see methods section). It also had an impact in the results presented below.

### Optimization runtime profiles

The DE’s optimization runtime profiles (mean fitness and SD of fitness) for the three datasets (simulated cattle, simulated pig, and real pig) and the two DL models (MLP and CNN) are shown in Figures 5-7. The mean fitness increased during the DE run, but it is important to note that it can - and did - also decrease at some points due to the stochastic sampling of individual subsets that we used for model testing to avoid overfitting (panels A and C of Figures 5-7). A similar short-term decrease in fitness was observed when using DE to optimize model hyperparameters in the context of emotion recognition (Nakisa *et al.* 2018). In our case, the occasional drop in the mean fitness was due to the retraining of the models as the refitting of the same model could yield a lower fitness. Thus, sometimes, even if a current *Titleholder* won the challenge, its new fitness could be lower than before due to the re-fitting. Alternatively, when a new *Challenger* won the contest, its fitness could have been higher that the refitted fitness of the *Titleholder* but still lower than the previously estimated fitness for that *Titleholder,* which resulted in a new candidate solution in the population but also in a lower fitness.

**Figure 5.**
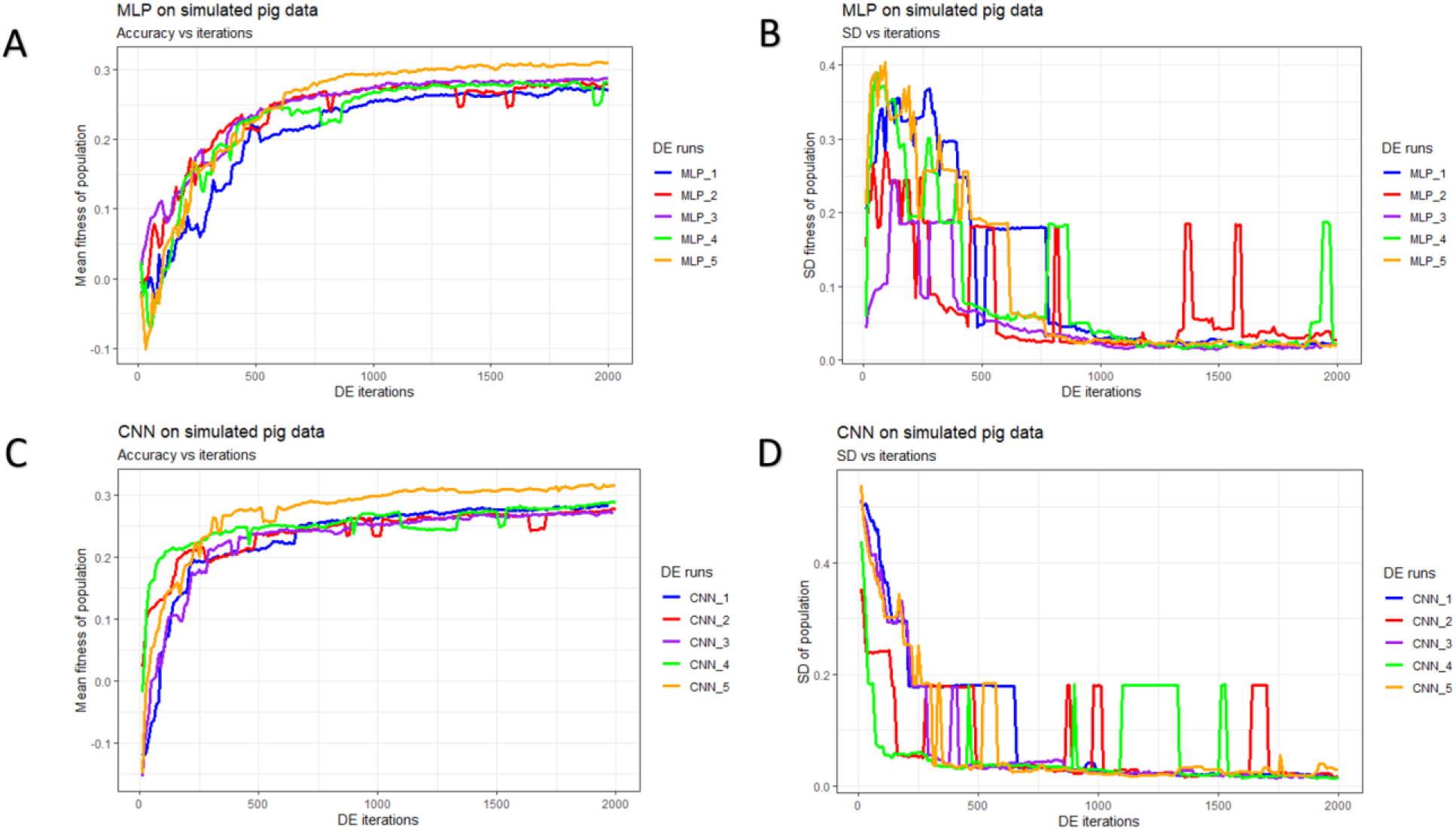
History of differential evolution by algorithm and data partition in the simulated pig dataset over 2,000 iterations. Mean and standard deviation of the fitness (correlation between the predicted and true phenotype) were computed given each population. (A) Mean fitness of five populations by fitting multilayer perceptron (MLP) models. (B) Standard deviation of fitness within each population (MLPs). (C) Mean fitness of five populations by fitting convolutional neural network (CNN) models. (D) Standard deviation of fitness within each population (CNNs).

**Figure 6.**
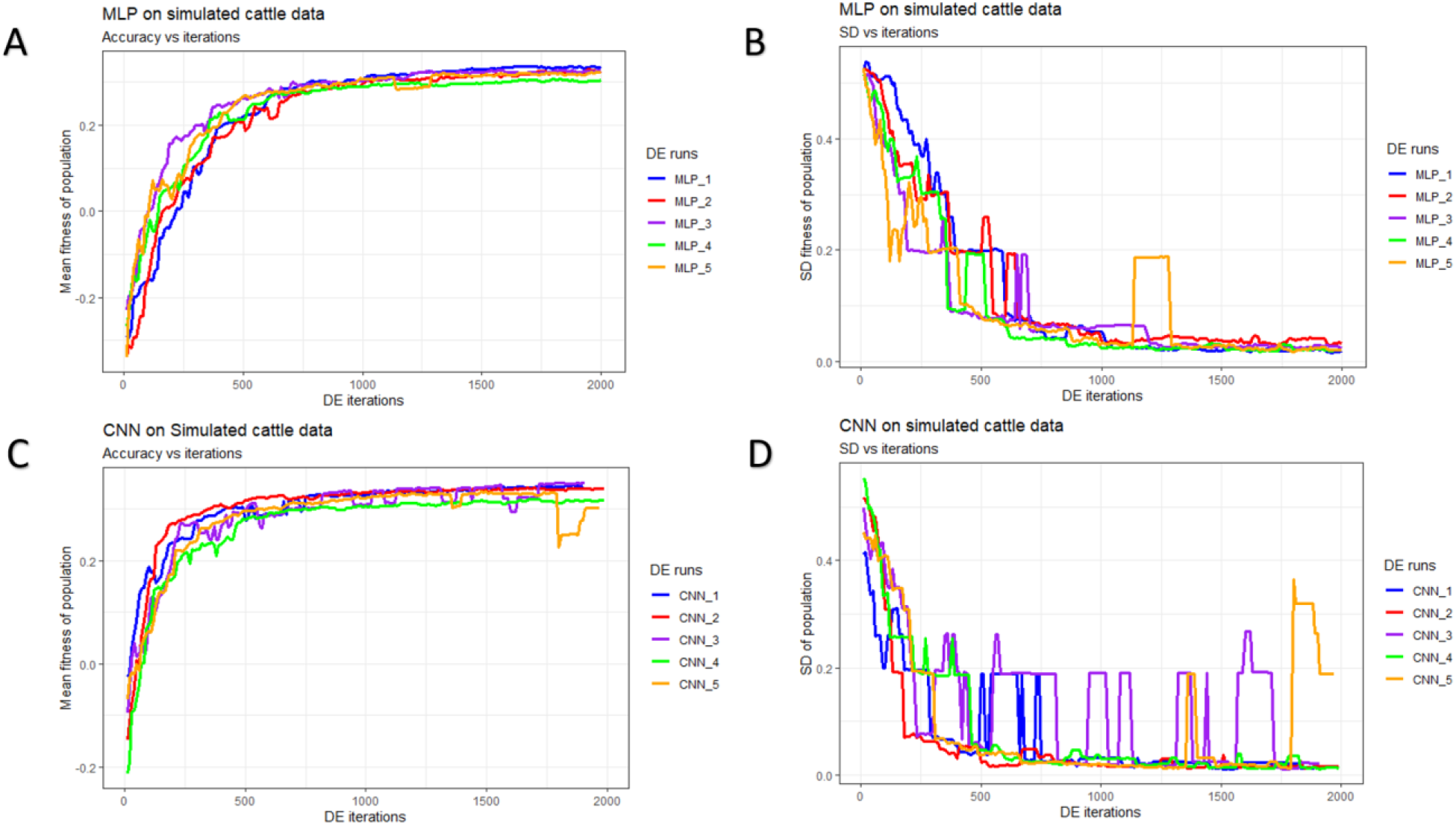
History of differential evolution by algorithm and data partition in the simulated cattle dataset over 2,000 iterations. Mean and standard deviation were computed given each population. (A) Mean fitness of five populations by fitting multilayer perceptron (MLP) models. (B) Standard deviation of fitness within each population (MLPs). (C) Mean fitness of five populations by fitting convolutional neural network (CNN). (D) Standard deviation of fitness within each population (CNNs).

**Figure 7.**
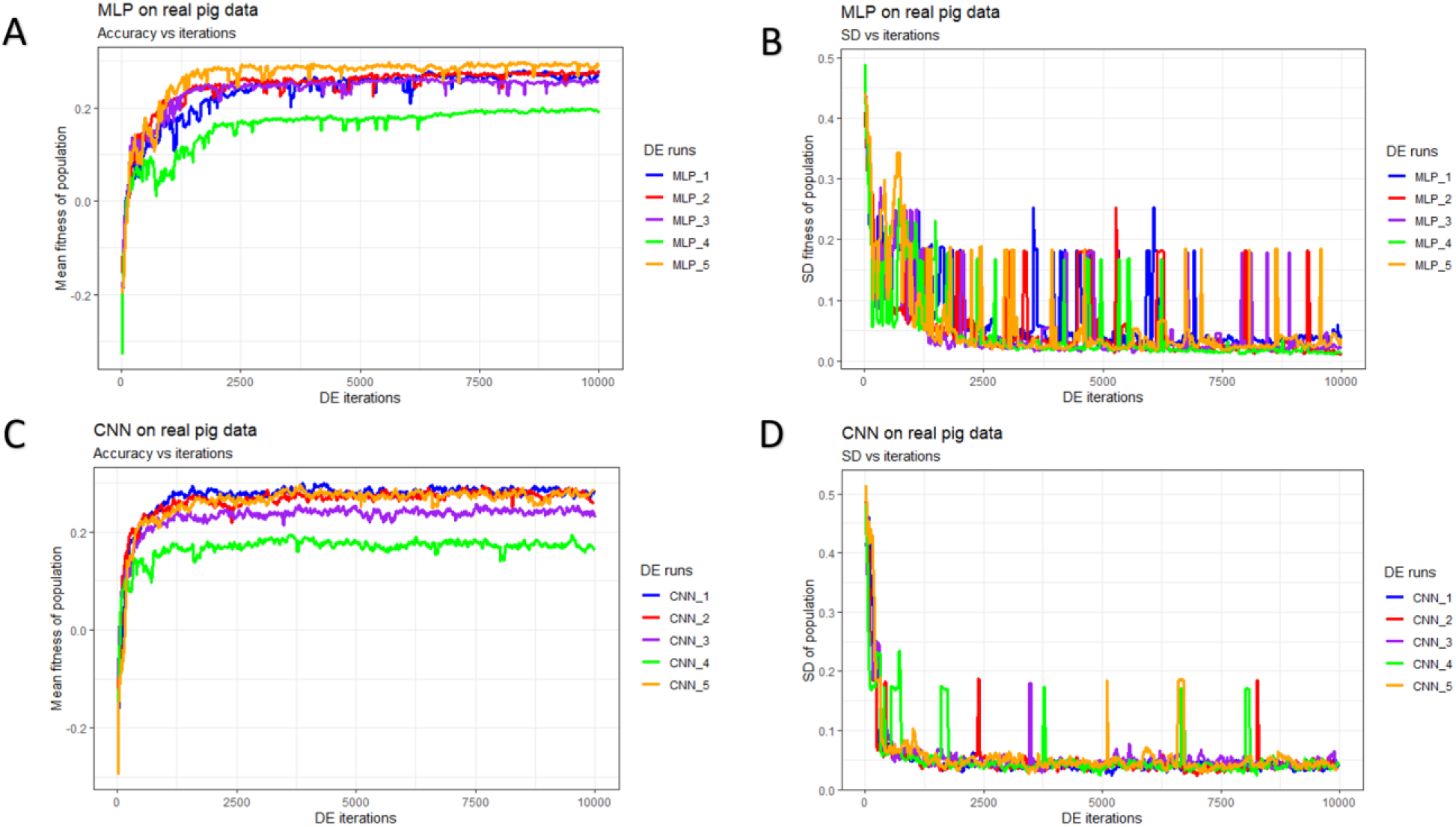
History of differential evolution by algorithm and data partition in the real pig dataset over 10,000 iterations. Mean and standard deviation were computed given each population. (A) Mean fitness of five populations by fitting multilayer perceptron (MLP) models. (B) Standard deviation of fitness within each population (MLPs). (C) Mean fitness of five populations by fitting convolutional neural network (CNN) models. (D) Standard deviation of fitness within each population (CNNs).

In general, DE for CNNs converged faster (reached the maximum possible average fitness) compared to DE for MLPs (panels A/C of Figures 5-7). For CNNs, DE converged after approximately 600, 700, and 1,500 iterations for the simulated pig, simulated cattle, and real pig datasets, respectively. For MLPs, DE converged in approximately 1,000, 1,000, and 2,500 iterations for the three datasets, respectively. One possible explanation is that, MLP disregards spatial information and use each neuron as an independent predictor, while CNN tends to learn from a global pattern at the beginning and then summarize the features into a local level (Lecun *et al.* 2015). In genomic data, linkage disequilibrium is a nonrandom relationship of alleles at different physical locations, which is a sensitive indicator that structures a genome (Slatkin 2008). Also, Tang and Sun (2019) argued that CNN could be utilized to extract motifs from homologous sequences, where motifs are essential features for distinguishing different sequence families. Given a dataset with spatial structure, CNN potentially has advantage over MLP that CNN can deal with local connectivity.

For each dataset, evolved MLPs and CNNs converged to similar mean fitness but varied across data partitions (panels A/C of Figures 5-7). The mean fitness in the simulated pig dataset ranged from 0.27 to 0.31 and the range was (0.31, 0.33) for the simulated cattle dataset, while the real pig dataset had a range of (0.19, 0.29). Mitchell et al. (2015) trained networks with permuted datasets and also reported varying predictive performance given different data partitions.

Most evolved populations had a fitness SD smaller than 0.05. However, one exception was population 5 with CNNs in the simulated cattle dataset (panel D of Figure 6). Only this population had a large SD of 0.19 and the population contained a CNN hyperparameter set with penalized fitness, indicating its failure to remove a penalized individual in 2,000 iterations. DE performance is sensitive to the number of iterations set by the user and generally solutions can evolve further when the iteration number is increased (Gämperle *et al.* 2002; Kok and Rajendran 2016). Thus, the solution with penalized fitness should be removed by introducing more DE iterations. On the other hand, the post-DE refitting (described in top model selection) would further exclude this solution. Overall, the within population SDs for both MLP and CNN models were reduced over DE iterations (panels B/D of Figures 5-7), suggesting evolved models in each population had similar performance. Kim and Lee (2019) reported that deep learning models with different hyperparameters could have the same predictive performance, which indicated that the best solution may not be unique. Zhang et al. (2020) also indicated that superior solutions would prefer the closest candidates in evolutionary optimization algorithms. Therefore, we argue that DE evolves a population to where candidate solutions are increasingly similar to each other. Furthermore, distributions of evolved models showed similarities in hyperparameter options e.g. activation function, number of layers, filter size, optimizer, dropout, and pooling, while the hyperparameters were less similar in number of nodes (filters), fully connected layer in CNN, batch size, and L2 regularization (Tables S4-S15). Yu and Zhu (2020) have mentioned that in the process of optimization, hyperparameters with greater importance received preferential treatment, whereas it was difficult to quantitatively determine the significance of the hyperparameters. We argue that the dissimilarity in the hyperparameters resort to the less important hyperparameters.

### Characteristics of selected hyperparameters

Table 2 shows the top MLPs from each population (one hyperparameter solution from each population, 15 in total). Activation functions of MLPs optimized for the simulated datasets varied in “elu”, “selu”, “relu”, “softplus” and “linear”, while in optimized MLPs for the real pig dataset, the “sigmoid” function was fixed across all selected individuals. Noteworthy: In this study, the input of the DL model was the allelic count of one of the alleles (coded as 0, 1 and 2), thus, all the input nodes were non-negative values. Interestingly, “elu”, “selu” and “relu” are almost identical when the input is a non-negative value, and the “linear” activation is very similar to those functions too (differing only in the slope). Moreover, “softplus” and “sigmoid” are the most different activation functions compared to the elu-linear family. The activation functions of the top models are described in File S1. Our finding agrees with Bellot et al. (2018) who suggested “elu”, “softplus” and “linear”, and also “relu” recommended by Pérez-Enciso and Zingaretti (2019). Moreover, as the simulated datasets were generated by only considering additive genetic effects, QTLs were contributing to the response linearly. Therefore, we speculate that the optimized DL models for the simulated datasets potentially unveiled the additive genetic effect by selecting predominantly the linear-like activation functions. However, if the input allelic counts were normalized, these same activation functions may produce different predictive performance, as the input could be a negative value, and over that range of inputs, elu, relu, selu and linear have very different output behaviors. For the real dataset, optimized MLPs fixed the non-linear activation function “sigmoid”. We argue that the selected non-linear activation reflects the increased complexity of real datasets e.g. allele interaction effects, such as dominance, that potentially exist in a real dataset. Regarding this perspective, Zingaretti et al. (2020) indicated that in a real dataset, DL could model complex relationships by employing non-linear functions, and they also observed that sigmoid-like hyperbolic tangent (tanh) was a safer choice overall. Models for the simulated datasets selected one-layer, two-layer, and three-layer MLPs, while all MLPs for the real pig data were three-layer models. MLPs differed in number of neurons, batch size, and epochs. The optimizers of selected MLPs for the simulated datasets focused on “adam” and “adamax”, while for the real dataset “sgd” was further included. Dropout rates of MLPs were between 0 and 0.034 for the simulated datasets and were between 0.182 and 0.617 for the real pig dataset. Compared to the model architectures selected by Bellot et al. (2018) and Pérez-Enciso and Zingaretti (2019), we had similar hyperparameter options in number of layers and activation function. But we selected different optimizers and the dropout in our case tended to be larger in the real pig dataset. Penalty weights for L2 regularization of MLPs had a range of (0.01, 0.16) for the simulated datasets and a range of (0.03, 0.85) for the real pig dataset. We did not find any suggested L2 weight applied to genomic prediction studies.

**Table 2.** Hyperparameters of selected MLP models from each population

| **Dataset** | **DE No.** | **activation** | **No. layer(s)** | **No. neurons** | **batch** | **epoch** | **optimizer** | **dropout** | **L2** |
| --- | --- | --- | --- | --- | --- | --- | --- | --- | --- |
| SP | 1 | elu | 2 | [446,87] | 51 | 37 | adam | 0.006 | 0.06 |
| SP | 2 | elu | 2 | [412,150] | 41 | 45 | adam | 0.020 | 0.16 |
| SP | 3 | elu | 2 | [470,155] | 46 | 44 | adam | 0.015 | 0.06 |
| SP | 4 | selu | 2 | [474,145] | 54 | 45 | adam | 0.032 | 0.13 |
| SP | 5 | softplus | 2 | [397,87] | 54 | 45 | adam | 0 | 0.13 |
| SC | 1 | elu | 3 | [429,330,57] | 44 | 28 | adam | 0.030 | 0.04 |
| SC | 2 | relu | 2 | [411,106] | 48 | 41 | adamax | 0.002 | 0.06 |
| SC | 3 | elu | 3 | [401,269,93] | 11 | 27 | adamax | 0.001 | 0.01 |
| SC | 4 | relu | 1 | 409 | 56 | 21 | adam | 0.034 | 0.14 |
| SC | 5 | relu | 1 | 444 | 47 | 33 | adam | 0.020 | 0.16 |
| RP | 1 | sigmoid | 3 | [374,192,25] | 10 | 40 | sgd | 0.352 | 0.85 |
| RP | 2 | sigmoid | 3 | [476,193,69] | 54 | 42 | adam | 0.480 | 0.52 |
| RP | 3 | sigmoid | 3 | [483,291,8] | 44 | 46 | adamax | 0.182 | 0.12 |
| RP | 4 | sigmoid | 3 | [457,234,79] | 31 | 41 | adamax | 0.465 | 0.03 |
| RP | 5 | sigmoid | 3 | [386,251,148] | 8 | 40 | sgd | 0.617 | 0.75 |

SP, simulated pig dataset; SC, simulated cattle dataset; RP, real pig dataset; DE No., differential evolution of different data partition; No. layer(s), number of hidden layers; No. neurons, number of neurons according to the number of hidden layers.

Table 3 shows top CNNs. Optimized CNNs had three options for activation function: “linear”, “elu”, and “relu” (File S1). Similar to our results, top CNNs selected by Bellot et al. (2018) also included “linear” and “elu”, while Pérez-Enciso and Zingaretti (2019) used “relu”. The number of convolutional layers varied from one to three while all CNNs for the simulated datasets fixed with one. Notably, Bellot et al. (2018) also selected one-layer and three-layer CNNs. Models differed in number of filters, epoch, and the number of neurons in the fully connected layer. The filter sizes tended to be larger in the selected models. The large filter sizes were different from other studies that suggested two or three (Bellot *et al.* 2018; Pérez-Enciso and Zingaretti 2019). Optimizers of selected CNNs were “adamax”, “rmsprop” and “adam” while CNNs for the real pig data fixed “adam”, and this finding is different from the “nadam” obtained by Pérez-Enciso and Zingaretti (2019). Most CNNs across the three datasets used average pooling for the pooling layer. For genomic prediction studies, we did not find a suggested pooling option in the literature. Dropout rates of CNNs ranged from 0.008 to 0.827 and the range was smaller (0.021,0.277) for the real pig dataset. However, our finding in dropout differed from the small dropout (5-10%) recommended by Pérez-Enciso and Zingaretti (2019). Most L2 penalty weights were smaller than 0.16 while there were three exceptions (0.52, 0.75 and 0.85).

**Table 3.** Hyperparameters of selected CNN models from each population

| Data | DE No. | activation | No. layers | No. filters | filter size | epoch | FCL | optimizer | dropout | L2 | Pooling |
| --- | --- | --- | --- | --- | --- | --- | --- | --- | --- | --- | --- |
| SP | 1 | linear | 1 | 110 | 19 | 25 | 17 | adamax | 0.197 | 0.21 | average |
| SP | 2 | elu | 1 | 16 | 15 | 32 | 110 | rmsprop | 0.146 | 0.03 | average |
| SP | 3 | elu | 1 | 15 | 8 | 44 | 79 | rmsprop | 0.692 | 0.02 | average |
| SP | 4 | linear | 1 | 59 | 20 | 24 | 49 | adamax | 0.496 | 0.23 | max |
| SP | 5 | linear | 1 | 109 | 13 | 27 | 109 | adam | 0.827 | 0.01 | average |
| SC | 1 | linear | 1 | 116 | 20 | 30 | 16 | adam | 0.370 | 0.10 | average |
| SC | 2 | linear | 1 | 87 | 12 | 25 | 12 | adam | 0.086 | 0.13 | average |
| SC | 3 | linear | 1 | 32 | 8 | 42 | 24 | adam | 0.250 | 0.19 | average |
| SC | 4 | linear | 1 | 79 | 20 | 44 | 27 | adamax | 0.666 | 0.06 | max |
| SC | 5 | linear | 1 | 98 | 16 | 40 | 153 | adam | 0.151 | 0.17 | average |
| RP | 1 | elu | 2 | [51,113] | 18 | 22 | 50 | adam | 0.277 | 0.67 | average |
| RP | 2 | relu | 3 | [24,81,121] | 12 | 27 | 268 | adam | 0.067 | 0.11 | average |
| RP | 3 | elu | 2 | [64,112] | 13 | 45 | 278 | adam | 0.021 | 0.87 | average |
| RP | 4 | relu | 3 | [44,73,106] | 13 | 47 | 326 | adam | 0.008 | 0.18 | average |
| RP | 5 | elu | 3 | [41,71,128] | 5 | 41 | 238 | adam | 0.051 | 0.35 | average |

SP, simulated pig dataset; SC, simulated cattle dataset; RP, real pig dataset; DE No., differential evolution of different data partitions; No. layers, number of convolutional layers; No. filters, number of filters applied based on No. layers; FCL, size (number of neurons) of the fully connected layer after flatten layer.

Despite our evolved hyperparameter sets being similar to those described in the literature (Bellot *et al.* 2018; Pérez-Enciso and Zingaretti 2019), part of hyperparameter configurations e.g. number of nodes (filters), optimizer, and dropout differed from those described in the existing studies. This is likely due to that the optimal hyperparameter configuration depends on the specific genomic dataset, and a hyperparameter’s relevance may depend on another hyperparameter’s value (Luo 2016). As Bellot et al. (2018) worked on a human dataset and Pérez-Enciso and Zingaretti (2019) investigated a wheat dataset, we attribute the variation among optimized hyperparameters to the specific dataset. It is also possible that our extended hyperparameter space searched for more instances, which led to the differences in some hyperparameters compared to other studies. While other researchers optimized hyperparameters by discretizing the parameter space, we regarded number of neurons (filters), dropout, L2 regularization, batch size, epoch, and filter size as continuous values, which considerably expanded the hyperparameters search space.

### Performance of optimized models under cross validation

The objective of this paper is to provide a framework to optimize DL hyperparameters for genomic prediction and not to compare the optimized DL with GBLUP. It is however still relevant to use GBLUP as a baseline of reference prediction methods to contextualize our results (see Figure S1). For the simulated datasets, GBLUP was the slightly better than the rest of the models. A similar result was presented by Abdollahi-Arpanahi et al (2020). This is not surprising in our study because GBLUP (described in File S1) is a model well suited for the simulated data which is entirely additive and composed of a large number of very small effects that approximate the infinitesimal model. However, for the real pig dataset, the pattern was somewhat different, and the best performing model was dependent on the data partition. As explained later in this section, we attribute this phenomenon to the small sample size of the real pig dataset. D’souza et al. (2020) argued that for a small dataset (e.g.: N<5000), the presence of substructure or even a few outliers may have a profound influence on the predictive performance under a specific data partition, skewing the overall estimate of the predictive performance and affecting the outcome of any optimization method that is used.

As DL is a methodology that relies on a learning process conditioned on the problem that it is solving (Montesinos-López *et al.* 2018), it is less likely that a DL model can achieve its best possible prediction accuracy using a hyperparameter set optimized from other independent studies. To investigate this, we trained MLPs and CNNs with hyperparameters selected for predicting human traits (Bellot *et al.* 2018) and for a wheat dataset (Pérez-Enciso and Zingaretti 2019), across the three datasets in this study. Table S16 shows hyperparameters of MLPs and CNNs obtained from the two studies. Figures 8-10 shows the predictive performance of random DL models, optimized DL models, and top DL models selected by Pérez-Enciso and Zingaretti (2019). These models were applied to all three datasets. Randomly selected DL models and optimized DL models differed in training data partitions due to independent DE optimizations performed within each partition, while the models suggested by the two previous studies (Bellot *et al.* 2018; Pérez-Enciso and Zingaretti 2019) were fixed in all partitions. Average prediction accuracies of (external) cross validations and SD of the correlations were obtained by refitting each model 30 times. The panels in Figures 8-10 represent the predictive performance of competing DL models for each data partition within each dataset. Noteworthy, the optimized models using DE were consistently the best when compared to randomly chosen models or to models taken from the literature, that have been optimized for other datasets.

**Figure 8.**
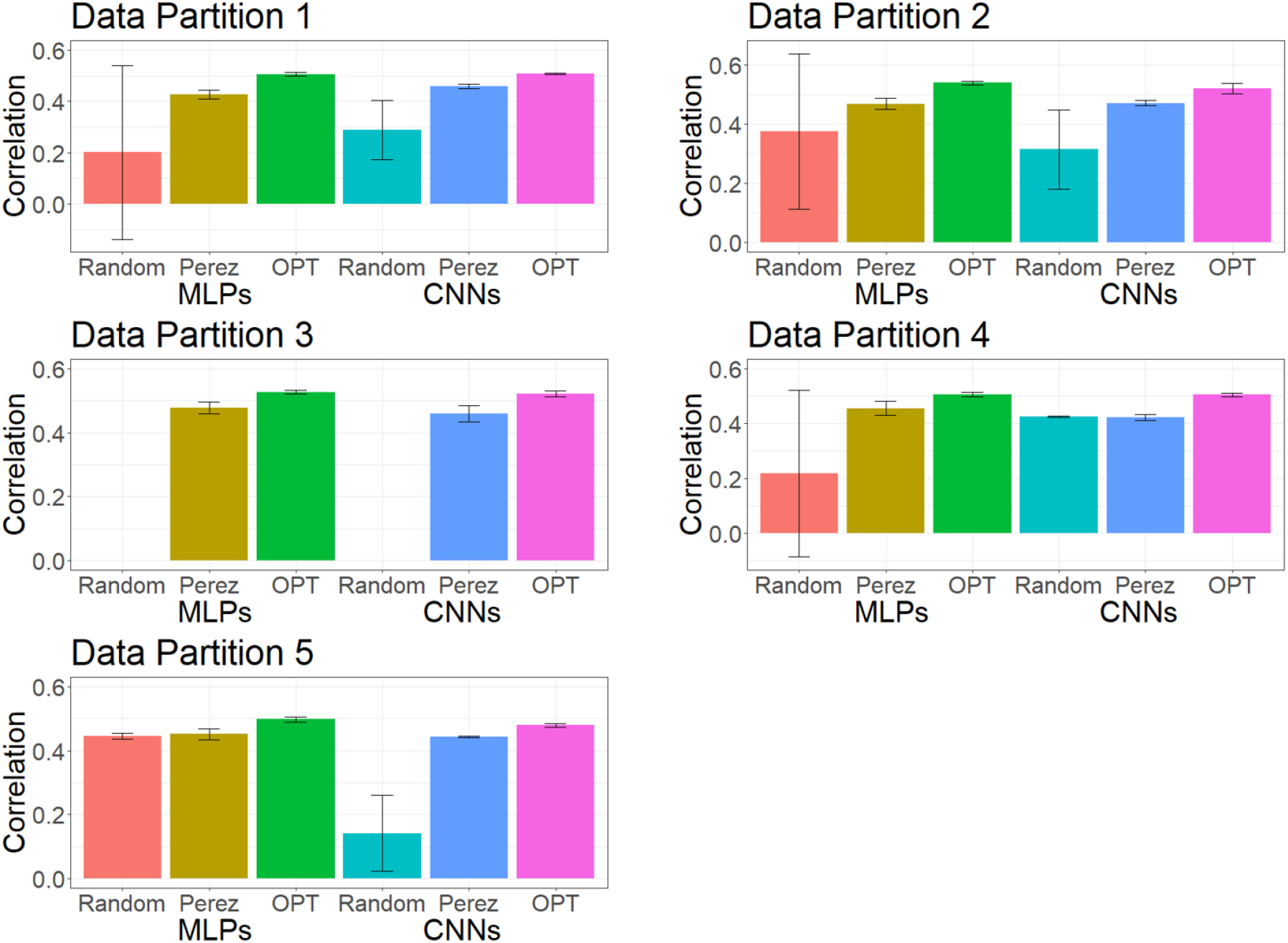
Mean predictive performance and error bars of MLPs and CNNs using different hyperparameters. Models were tested on five data partitions of the simulated pig dataset. The error bar represents the mean ± standard deviation of cross validation by fitting the same model 30 times. The left three bars are MLP models and the right three bars are CNN models. Null bar means the model did not converge. Random, random hyperparameters; Perez, hyperparameters recommended by Pérez-Enciso and Zingaretti (2019); OPT, optimized hyperparameters using DE. Abbreviations stand for the same meaning in Figures 9 and 10.

Models with hyperparameters chosen by Bellot et al. (2018) did not converge in any data partition and so they are not shown in Figures 8-10. This was likely due to exploding gradients or vanishing gradients (as previously discussed). Another observed case was that the model predicted every individual with the same value, making it impossible to compute the correlation between the predicted and observed response. This also confirmed the observation by Bellot et al. (2018) that convergence problems persisted after reinitializations of the algorithm.

For the simulated pig dataset, the MLP and the CNN suggested by Pérez-Enciso and Zingaretti (2019) was slightly worse than the optimized MLPs and CNNs that we obtained with DE. However, their performance was much worse in the simulated cattle and the real pig datasets. Again, the optimal hyperparameter configuration is problem-dependent and thus, it is important to search for the proper hyperparameters in DL genomic prediction applications given a specific dataset.

The variations in the predictive performance under re-training observed in all models indicated that DL models were likely overfitting the data. Abdollahi-Arpanahi et al. (2020) showed variance in predictive performance (in terms of accuracy and mean squared error) of MLPs and CNNs over 10 replicates of cross validation which is in agreement with our results. In general, the SD of the correlation between predicted and observed phenotypes for the optimized MLPs/CNNs and those proposed by Pérez-Enciso and Zingaretti (2019) were smaller in the simulated datasets, while the SD in the real pig dataset was larger (Figure 10 compared to Figures 8 and 9). We speculate that there are two possible reasons for the variation: DL models are initialized with random weights at starting points and a relatively small sample size for training. For the random weights at baseline, Bellot et al. (2018) explained that the performance of MLPs and CNNs depended on initialization values. For the training sample size, Abdollahi-Arpanahi et al. (2020) indicated that larger sample sizes improved the predictive ability of DL methods. Furthermore, in the field of image classification, Shahinfar et al. (2020) showed increased prediction accuracy and reduced variation in the performance of DL models as the sample size grew. Based on the results in Figures 8-10, the merit in terms of lower SD over replicates of cross validations was clearer in the simulated datasets that had larger sample sizes (N=7,539 for both the simulated pig and the simulated cattle datasets). In the real pig dataset that had a smaller sample size (N=910), SD was larger compared to those in the simulated datasets. Therefore, we argue that both the predictive ability and variation in the same DL models are associated with training sample size. Montesinos-López et al.(2018) also mentioned that DL method may fail to learn a proper generalization of the knowledge contained in the data, given small datasets.

**Figure 9.**
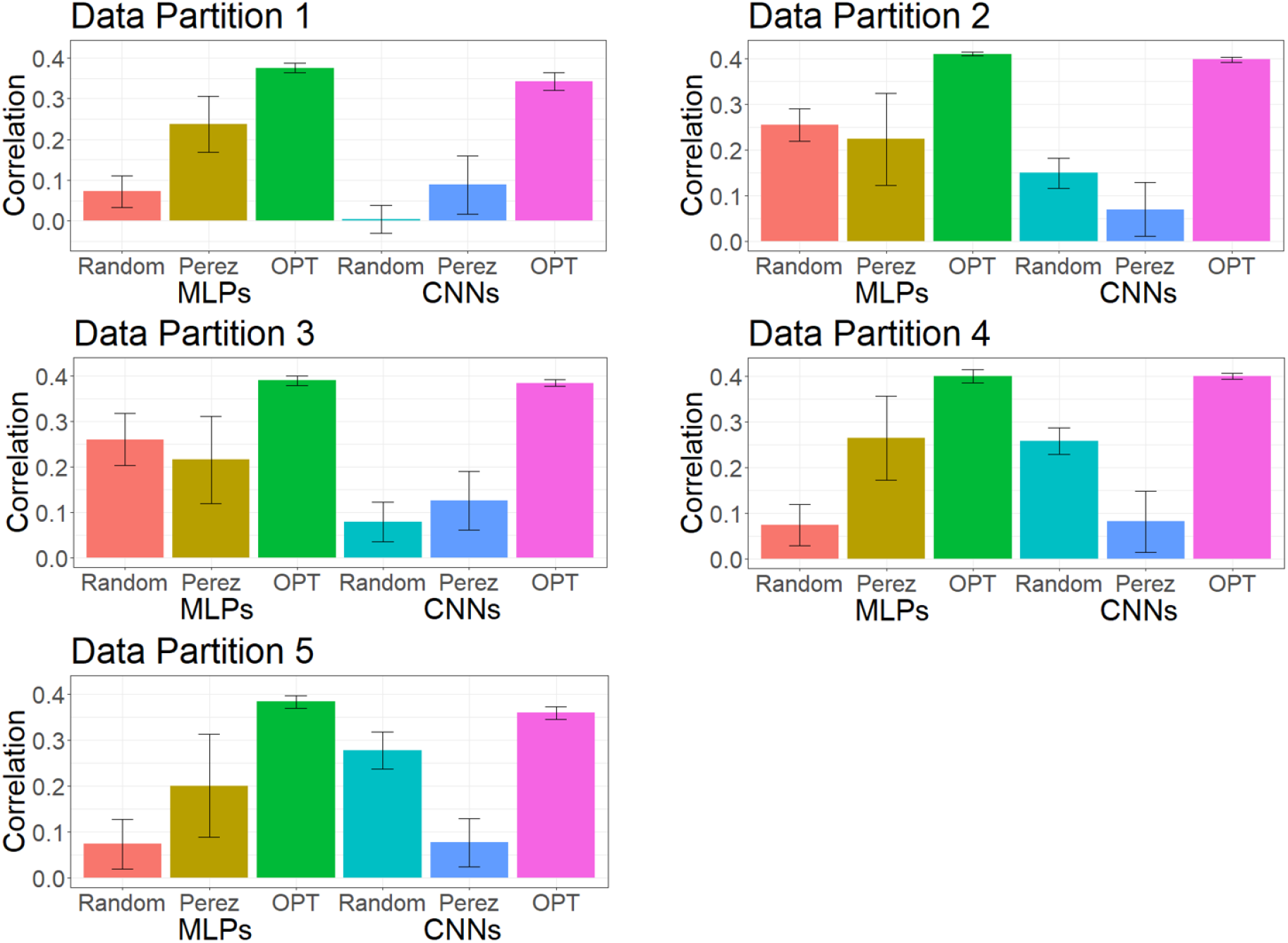
Mean predictive performance and error bars of MLPs and CNNs using different hyperparameters. Models were tested on five data partitions of the simulated cattle dataset. The error bar represents the mean ± standard deviation of cross validation by fitting the same model 30 times. The left three bars are MLP models and the right three bars are CNN models.

**Figure 10.**
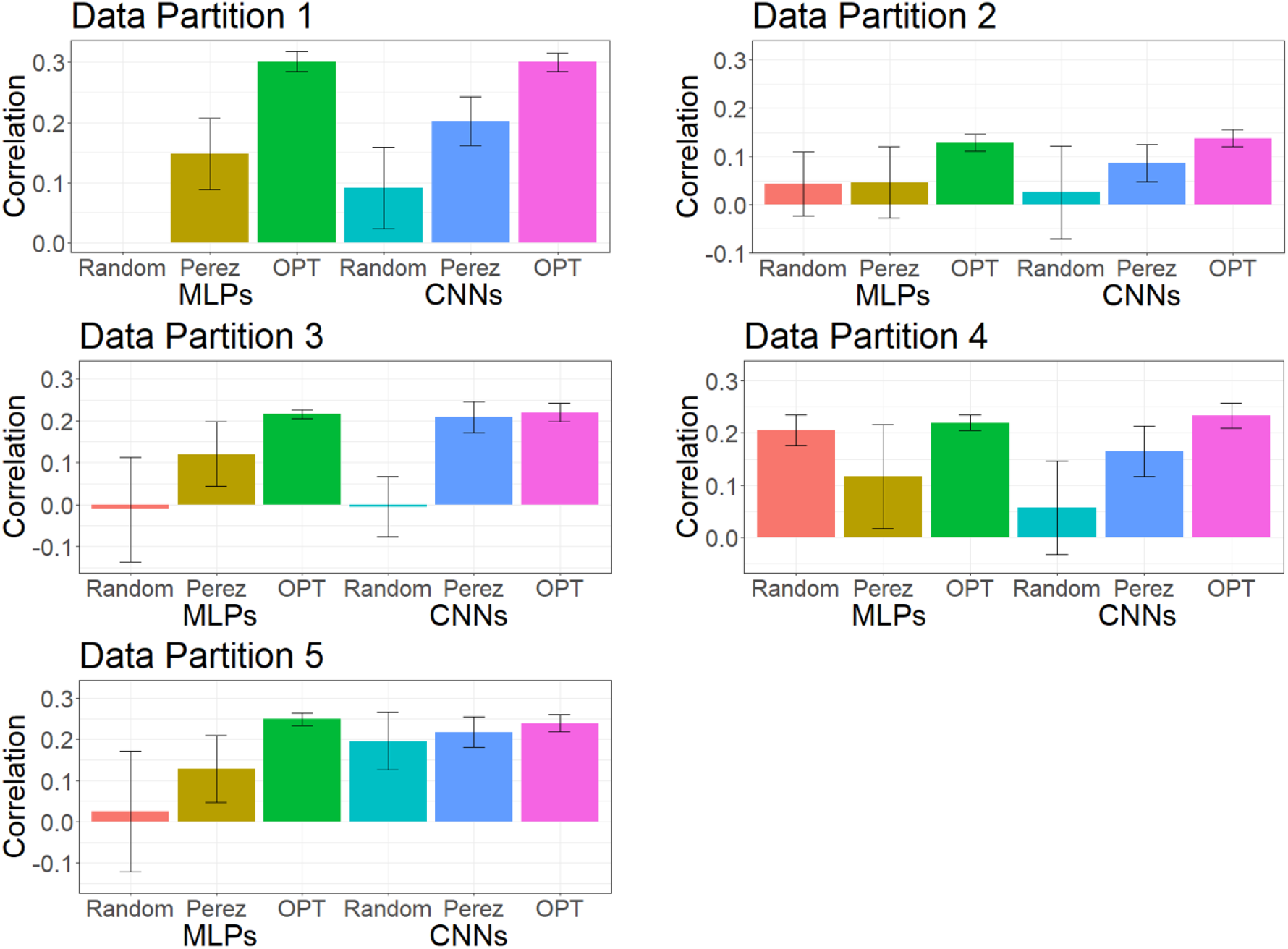
Mean predictive performance and error bars of MLPs and CNNs using different hyperparameters. Models were tested on five data partitions of the real pig dataset. The error bar represents the mean ± standard deviation of cross validation by fitting the same model 30 times. The left three bars are MLP models and the right three bars are CNN models. Null bar means the model did not converge.

## Conclusions

Overall, DL can be adapted to perform genomic prediction of complex traits, but it requires some effort to select appropriate hyperparameters. Any hyperparameter optimization will likely be dataset-specific and characteristics such as population structure and genetic architecture of the predicted trait may well require different DL model hyperparameters. In this study, we implemented differential evolution (DE) as a method to simultaneously identify optimal combinations of multiple hyperparameters. Compared to randomly selected models, our optimized MLPs and CNNs showed significant improvement in the predictive performance. In comparison to DL models with hyperparameters selected from other studies, optimized MLPs and CNNs also yielded better predictive accuracy. DE is an efficient and semi-automatic algorithm that can be used to select an optimal hyperparameter set that leads to a better predictive performance. Moreover, overparameterization of DL can be mitigated by refitting models and selecting those that produce more consistent (less variable) prediction accuracies. We showed that this is more important when working with small datasets.

## Acknowledgements

This work was supported by Agriculture and Food Research Initiative Awards No. 2017-67007-26176, No. 2010-65205-20342, the National Institute of Food and Agriculture (AFRI Project No. 2019-67015-29323), and by funding from the National Pork Board Grant No. 11-042. Partial funding was also provided by the US Pig Genome Coordinator.

